# VirNA: a novel Minimum Spanning Networks algorithm for investigating viral evolution

**DOI:** 10.1101/2024.11.05.620028

**Authors:** G. Mazzotti, L. Bianco, M. Bado, E. Lavezzo, S. Toppo, P. Fontana

**Author notes:** Authors to whom correspondence should be addressed Stefano Toppo, Università degli Studi di Padova, Dipartimento di Medicina Molecolare, Paolo Fontana, Fondazione Edmund Mach. These authors contributed equally (S. Toppo, P. Fontana).

## Abstract

Next Generation Sequencing technologies are essential in public health surveillance for tracking pathogen evolution, spread, and the emergence of new variants. However, the extensive sequencing of viral genomes during recent pandemics has highlighted the limitations of traditional molecular phylogenetic algorithms in capturing fine-grained evolutionary details, emphasizing the urge for more effective approaches to manage these large-scale data. VirNA (Viral Network Analyzer) addresses this challenge by reconstructing detailed mutation patterns and tracing pathogen evolutionary routes in specific regions through Minimum Spanning Networks. It analyzes thousands of sequences, generating networks where nodes represent identical genomic sequences linked to their metadata, while edges represent evolutionary pathways.

**AUTHOR SUMMARY:** The authors present Viral Network Analyzer (VirNA), a new tool designed to help track the spread of viruses during pandemics. During recent outbreaks, researchers faced challenges due to the sheer volume of viral genome data. Traditional tools struggled to process this massive amount of information effectively. VirNA solves this problem by using a powerful method called Minimum Spanning Networks (MSN) with enhanced features to analyze large datasets quickly and accurately. VirNA allows scientists to map how viruses spread in specific areas, providing critical insights that other tools couldn’t achieve. It complements traditional phylogenetic methods, offering a detailed look at viral transmission routes. This makes it a valuable asset for public health surveillance, helping experts understand and respond to pandemics more effectively. Already tested in previous studies, VirNA represents a significant breakthrough for future pandemic preparedness, offering new ways to handle and interpret big data in viral research.

## 1. INTRODUCTION

The integration of Next Generation Sequencing technologies into pandemic surveillance has generated vast amounts of data, posing challenges for sequence management and interpretation. These data are essential for tracking emerging variants and reconstructing viral transmission routes. Recent global threats like SARS-CoV-2 and Human Monkeypox Virus have accelerated whole genome sequencing, generating large genomic datasets, and reshaping our understanding of viral evolution.

Molecular phylogeny is widely used to study pathogen evolution, but traditional phylogenetic algorithms struggle with rapidly evolving pathogens, often producing low-confidence trees due to the high similarity of viral sequences(3). To address this issue, we developed VirNA (Viral Network Analyzer), which uses Minimum Spanning Networks (MSNs)(4) to trace evolutionary pathways. VirNA can efficiently analyze thousands of viral genomes, integrating sampling dates and location metadata. This tool offers a new algorithmic strategy with respect to the existing MSNs tools Pegas(5) and PopArt(6) by integrating a new sequence connection criterion and enabling directional linkage between sequences.

To validate the performance of VirNA, we developed a benchmark and compared the results of VirNA to those obtained from PopArt(6) and Pegas(5), highlighting their key algorithmic differences. Additionally, we applied VirNA to datasets of real viral sequences and compared its output to that of phylogeny algorithms showing their complementary nature.

## 2. RESULTS AND DISCUSSION

### 2.1 VirNA

VirNA is an implementation of Bandelt et al.’s(4) with the introduction of novel elements for the enhancement of viral genome evolution analysis. Consistently with the original algorithm(4), connections between nodes represent the shortest paths, but with the addition of a constraint: the set of genomic mutations in a parent node must be fully contained within those of its child node. From this point forward, we will refer to this constraint as Haplotype Compatibility (HC). The directed edges between nodes represent the accumulation of new mutations in the child node. As a result, the algorithm forms a Directed Acyclic Graph (DAG), with edges directed from source nodes (fewer mutations) to target nodes (more mutations), adding a novel element compared to Bandelt et al.’s algorithm(4). Additionally, unlike Bandelt et al.’s algorithm(4), VirNA may split data into multiple distinct networks due to the mutation compatibility constraint.

The conceptual framework of VirNA is shown in Figure 1. In this network, nodes represent identical genomic sequences while directed edges represent potential evolutionary relationships based on the mutation accumulation principle with the minimum number of mutational events between sequences. Node A (mutation set {m_1_, m_2_}) is not connected to node B (mutation set {m_1_, m_13_}), due to the absence of mutation m_2_ in B, which breaks the compatibility requirement. While this can pose a constraint in the algorithm, availability of sequence-associated metadata can enable the user to reconsider the connections traced in the network.

**Figure 1:**
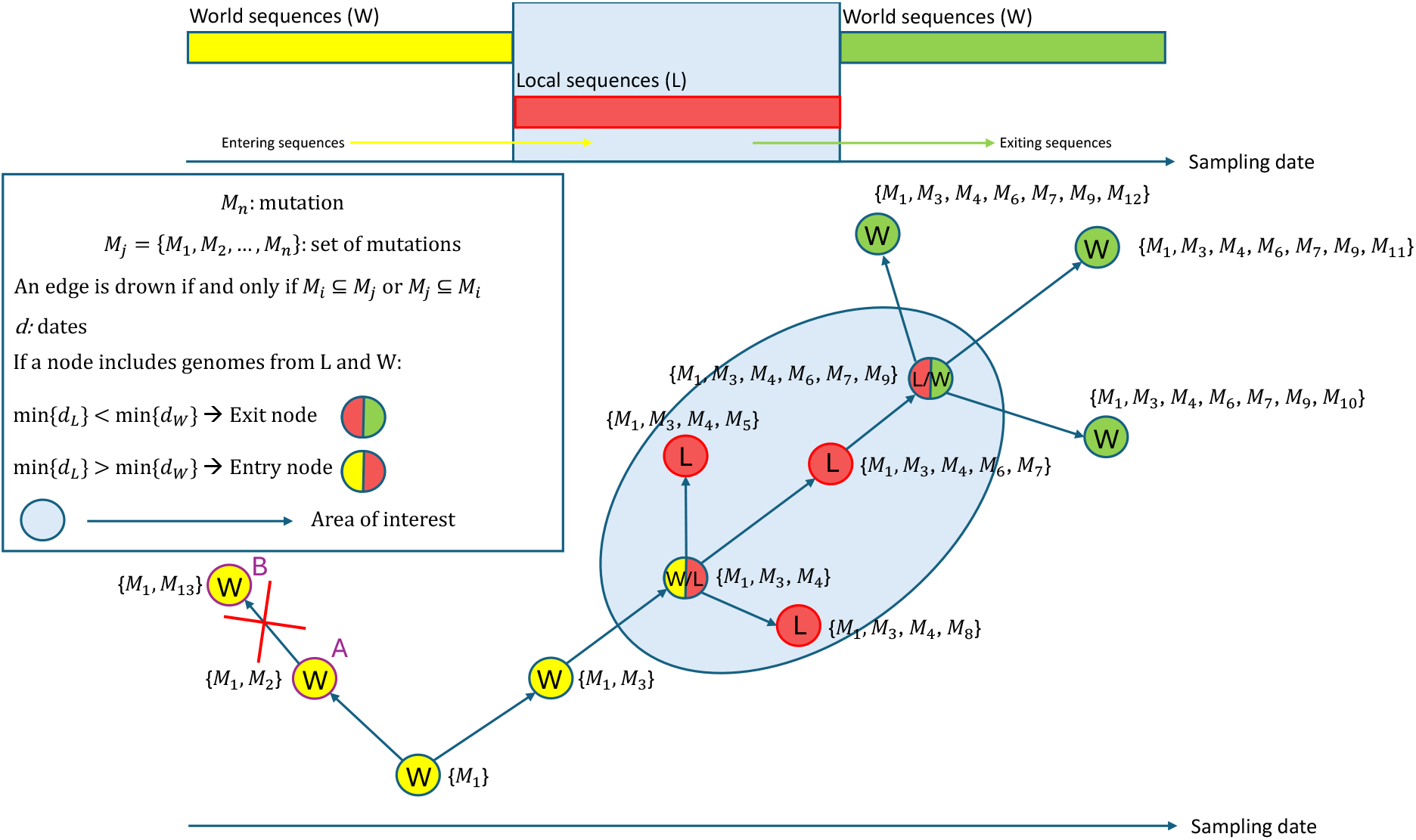
VirNA algorithm and the identification of “Entry” and “Exit” nodes. The figure depicts how VirNA algorithm works on a hypothetical example where twelve different viral genomic sequences are available. A connection between two nodes is established if and only if the mutations that define the target node precisely correspond to those characterizing the source node, complemented by additional mutations. VirNA is able to identify potential “entry” and “exit” nodes in the network. When a node in the network clusters both “Local” and “World” sequences (double colored nodes), if the earliest sampling date among the “Local” sequences is earlier than the earliest sampling date among the “World” sequences, the node is labeled as “exit” node (red and green node), conversely as “entry” node (red and yellow node).

VirNA also identifies potential viral introductions into or exits from specific geographical areas. We define the local area as “L” (“L” for “Local”, Figure 1) and all other regions as “W” (“W” for “World”, Figure 1). “Entry” nodes mark pathogen imports from outside the local area, while “exit” nodes indicate potential spread to other regions. If the earliest “Local” sequence predates the earliest “World” sequence in a mixed node, it is labeled as an “exit” node; otherwise, it is an “entry” node. Figure 1 illustrates an example: double-colored nodes represent sequences from both “Local” and “World” regions. The timeline indicates that local sequences were sampled middle period, with global sequences sampled earlier or later. The red-yellow node represents an entry point where local sequences followed world sequences, while the green-yellow node is an exit point with local sequences sampled earlier. An initial version of VirNA was employed to identify viral haplotypes entering and leaving Trento and Vo’ areas (Italy) during the pandemic(12,13).

### 2.2 Comparison with state of the art tools

The comparative analysis of the networks generated by PoPArt(6), Pegas(5), and VirNA confirms distinct topological structures arising from the different algorithms implemented by the tools. Both PoPArt(6) and Pegas(5) implement the Minimum Spanning Networks algorithm as proposed by Bandelt et al.(4), resulting in fully connected, undirected networks. VirNA, in addition, requires Haplotype Compatibility (HC) and directional connections, which lead to Directed Acyclic Graphs (DAGs) that follow a mutation accumulation rationale and may lead to split networks. We have examined the Haplotype Compatibility (HC) of the three tools and assessed how divergent the connections between nodes could become without applying this constraint, potentially leading to implausible networks. In our datasets, Pegas(5) identified over 60% of non-compatible connections (64% in the 1000-sequence dataset and 69% in the 2000-sequence dataset), while PoPArt(6) identified 9% and 6%, respectively. Pegas(5) linked non-compatible sequences at an average distance of 6 in both datasets, compared to PoPArt(6) mean distance of 3.6 (1000-sequences) and 3 (2000-sequences). Pegas(5) connected less than 12% of non-compatible sequences at a distance of 2, with most connections occurring at the average distance, while PoPArt(6) linked 34% (1000-sequences) and 40% (2000-sequences) at this shorter distance. Pegas(5) connected 13% of non-compatible sequences in the 2000-sequence dataset at a distance greater than 10, and 7% in the 1000-sequence dataset, while PoPArt(6) 3% and 4% respectively. We found out a link between two groups of sequences with entirely distinct mutation profiles:

○ Node one: seven mutations {‘A2480G’,‘C2558T’,‘G11083T’,‘G18255T’,‘C20428T’,‘C22088T’,‘G26144T’}
○ Node two: ten mutations {‘C1059T’,‘A1808G’,‘G4802A’,‘C5183T’,‘C14408T’,‘C18568T’,‘A23403G’,‘G24368T’,‘G25563T’,‘C26461T’}

Despite their high divergence, these sequences were sampled within a 3-day span (April 8–10, 2020) and collected by the same laboratory in the UK, making the connection between them unlikely. The compatibility criterion is likely to favor connections between sequences that have evolved from one another over a short evolutionary timespan. In the context of rapidly evolving pathogens, this leads to more reliable and accurate network reconstructions.

### 2.3 Comparison with phylogeny results

The comparison between the output topologies of Minimum Spanning Networks and Phylogenetic approaches confirms their strong complementary nature. VirNA effectively describes the evolutionary relationships among closely related sequences in both SARS-CoV-2 and Monkeypox case studies and distinguishes co-circulating non-compatible groups of sequences. In contrast, while phylogenetic analysis cannot detail the relationships among highly similar sequences, it provides insights into the broader evolutionary connections between the distinct groups of non-compatible sequences, being able to reconstruct the elements of the evolutionary history when dealing with distinct genomic sequences. All the details of the comparative analysis can be found in the Supplementary Materials.

### 2.4 Performance test of VirNA

A performance test was carried out on a server equipped with a 512 GB System memory and two AMD EPYC 32-Core Processors. VirNA was compared with PopArt(6) and Pegas(5) focusing on computational time. Five sets of SARS-CoV-2 sequences were tested, described in the Supplementary Materials and available on the GitHub page of VirNA. VirNA is always faster than the other two tools, as shown in Supplementary Table 1 and is the only tool able to manage large datasets containing over 10000 sequences.

## 3. MATERIALS AND METHODS

### 3.1 VirNA Algorithm Details

The software is implemented in Python(1) and Cython(2), exploiting the Numpy(7) library to speed up the calculation of the Hamming distance matrix, while igraph(8) is used for an efficient construction of the MSN. A Graph Modeling Language (GML) file is written and can be easily imported into Cytoscape(9), Gephi(10) or other similar tools for visualization or downstream analyses. VirNA requires a multiple alignment in FASTA format as input file. VirNA algorithm details and definitions are reported below

#### 3.1.1 VirNA Algorithm Definitions

○ Root sequence: the first sequence of the input multiple alignment is designated as the root sequence R and will not be included in the output network. This is the reference sequence against which mutations in the dataset are identified.
○ Input genomic sequences: let S={s_1_,s_2_,…,s_n_} be the set of input genomic sequences.
○ Mutation sets: for each sequence s_i_ where i=1,2,…,n, we define a set of mutations M_j_={m_1_,m_2_,…,m_n_}, representing the single-character differences between the root sequence R and the sequence s_i_.
○ Network nodes: identical sequences are grouped in a single node. Let G_k_={s_1_,s_2_,…,s_m_}, with m≤n, be the sets of nodes in the network, where s_1_=s_2_=_…_= s_m_.
○ Compatibility: The mutation set M_i_ is said to be compatible with the mutation set M_j_ if and only if M_i_⊂M_j_ or M_j_⊂M_i_.
○ Connected component: a connected component of a directed graph is defined as a subgraph where, for each pair of nodes, G_i_,G_j_ there either is a path from G_i_ to G_j_ or from G_j_ to G_i_;

#### 3.1.2 Initialization

○ For each node G_i_, compute the corresponding set of mutations M_i_ compared to the root R;
○ Calculate the Hamming distance among all sequences and store it in a list HD, HD=[HD_1,2_,HD_1,3_,…,HD_n-1,n_] where HD_i,j_(G_i_,G_j_) is the Hamming distance between nodes G_i_ and G_j_, with i: 1,…,n, j:i,…,n and i≠j;
○ Sort the list of Hamming distances HD in ascending order removing the duplicated values: SHD={HD_1_,…,HD_k_} where HD_i_ in HD, k<=n(n-1), HD_i_≠HD_j_ for each i≠j, HD_a_<HD_b_ for each a,b=1,…,k if a<b;
○ Define the root node G_R_ as the sequence with the lowest number of mutations compared to the reference R;
○ Create a starting network, MSN(0), where all the (grouped) sequences (nodes) are not connected;

#### 3.1.3 Iterative steps

At each step m>=1, let HD_m_ be the m-th distance in the sorted Hamming distances (SDH) list defined above. If no stop criteria are met (see below) the network MSN(m) is built from MSN(m-1) by adding all the edges from G_i_ to G_j_ if and only if all the conditions below are met:

○ HD_i,j_==HD_m_;
○ G_i_ is compatible with G_j_;
○ G_i_ and G_j_ do not belong to the same connected component;

Stop criteria at step m:

○ All the nodes G_i_ 1<=i<=n, are part of a single connected component of the current network MSN(m-1);
○ HD_m_>D, where HD_m_ is the m-th distance in SDH and D is a user-defined maximum allowable distance;

#### 3.1.4 Output

○ The final network represents the Minimum Hamming Distances among all input sequences;

### 3.2 Benchmark

An inhouse Python(1) script has been developed to assess the output of VirNA and compare it to the output networks generated by PoPArt(6) and Pegas(5). The script takes as input the aligned set of sequences S_i_={s_1_,s_2_,…,s_n_.} that the user wants to analyse using MSNs. For each sequence, the script generates a list of compatible sequences and cross-references it with the connections identified by VirNA to validate the accuracy of the traced links. Subsequently, the script compares the results produced by VirNA, PoPArt(6) and Pegas(5) to highlight the algorithmic differences among the tools. Specifically, the benchmark evaluates the networks generated by the tools, categorizing connections into three groups: those that are identical across all tools, those identified by VirNA but not by PoPArt(6) or Pegas(5), and those identified by PoPArt(6) or Pegas(5) but not by VirNA.

### 3.3 Real data and Phylogeny

We applied VirNA, along with Maximum Likelihood and Maximum Parsimony phylogenetic algorithms, to two real sequence datasets from the GISAID repository(11), comprising SARS-CoV-2 and Monkeypox sequences, to compare the outputs of these algorithmic approaches. Details on the datasets, as well as the phylogenetic tools and parameters used, are provided in the Supplementary Materials. Both the sets of sequences are available on the GitHub page of VirNA (https://github.com/font71/VirNA).

## 4. CONCLUDING REMARKS

VirNA is specifically designed to enhance the analysis and monitoring of rapidly evolving pathogens in large, fine-grained datasets, making it a powerful tool for pathogen surveillance. Unlike traditional MSNs(4), VirNA introduces a haplotype compatibility criterion and directional connections among nodes which reflect the accumulation of mutations, ensuring that only the most closely related sequences are linked. This approach produces evolutionary plausible networks, particularly suited for fast-evolving pathogens. By applying the compatibility criterion, VirNA enables users to clearly differentiate sequences with incompatible haplotypes that are co-circulating within the same time period, generating distinct networks whose evolutionary relationship can be explored through phylogenetic analysis. VirNA excels in datasets with minimal or no missing data, fully unlocking its potential in simulated data studies.

## 5. DATA AVAILABILITY

The findings of this study are also based on SARS-CoV-2 sequences and associated metadata available on GISAID and accessible at <u>10.55876/gis8.240212ht</u> and on MonkeyPox sequences and associated metadata available on GISAID (acknowledgements table available in the supplementary materials).

## 6. ACKNOWLEDGMENTS

We gratefully acknowledge all data contributors, i.e., the Authors and their Originating laboratories responsible for obtaining the specimens, and their Submitting laboratories for generating the genetic sequence and metadata and sharing via the GISAID Initiative, on which this research is based.

