## Supplementary Materials for "VirNA: a novel Minimum Spanning Networks algorithm for investigating viral evolution"

§ These authors contributed equally

* Authors to whom correspondence should be addressed

### S1. SARS-CoV-2 CASE STUDY

Three sets of SARS-CoV-2 genomic sequences, obtained from different timeframes and sourced from the EpiCov GISAID repository(1), were selected. All the sequences were collected across the United Kingdom and we decided to work with the Nexststrain nomenclature(2):

- Period 1: from October 16, 2021 to October 18, 2021. The Nextstrain clades(2) 21I (Delta) and 21J (Delta) are dominant in the UK.
- Period 2: from December 18, 2021 to December 22, 2021. The Nextstrain clades(2) 21J (Delta) and 21I (Delta) are still dominant across the UK but the clade 21K (Omicron) appears.

- Period 3: from May 09, 2022 to May 11, 2022. The Nextstrain clades(2) 21L (Omicron), 22A (Omicron), 22B (Omicron) and 22C (Omicron) are dominant in the UK. The Delta variants are no longer present.

The genomic sequences underwent a filtering process based on multiple criteria considering both genomic sequences and their associated metadata. The chosen sequences exhibit the following features: a complete collection date in YYYY-MM-DD format, a complete genome (>= 29000 b), absence of ambiguous characters, no head and/or tail gap trails or with gap trails shorter than 90 bases, and no insertions and/or deletions causing frameshift.

The cleaned genomic sequences were aligned in an all VS all fashion along with their reference sequences (NCBI Reference Sequence NC_045512.2) using the Nextstrain augur tool(3) with default parameters.

VirNA was applied to all the sequence sets, and only those sequences belonging to the largest hubs (comprising more than 4 members) were retained. This selection was made to emphasize the topological characteristics of the larger network hubs, while excluding isolated nodes.

We specifically selected sequences from the United Kingdom as a test set for observing the results of VirNA on real data, due to their abundance. After the cleaning steps, sequences from the following Nextstrain clades were retained: 21J (Delta) in period 1 (724 sequences), 21J (Delta) and 21K (Omicron) in period 2 (808 sequences) and 21L (Omicron) and 22C (Omicron) in period 3 (420 sequences). The ample availability of data allowed us to refine the initial dataset by applying stringent filtering criteria while maintaining its representativeness. To enhance the clarity and visually accessibility of the results, we selected specific time intervals, focusing on a limited number of sequences. These time periods were chosen to highlight the evolution of distinct viral variants in a definite geographic area.

The Minimum Spanning Networks generated by VirNA from the SARS-CoV-2 sequences are shown in Supplementary Figures 1 and 2, panels A, C and E. The background color of the networks, indicates the clade(2) to which the genomic sequences belong. The sequences labeled as 21J (Delta) have a light-yellow background and are present in both periods 1 and 2 (Supplementary Figures 1 and 2, panels A and E respectively). In period 2, the 21K (Omicron) sequences appear, marked with a light pink background. In period 3 (Supplementary Figures 1 and 2, panel E), the 21L (Omicron) sequences are backed by a light violet background, while 22C (Omicron) sequences are distinguished by a light green background.

VirNA distinguishes sequences with incompatible haplotypes that are co-circulating within the same time period. Incompatible sequences are those that do not meet the Haplotype Compatibility (HC) criterion, as outlined in the Materials and Methods section of the paper. In this case, the separated sequence groups correspond to distinct viral variants or sub-variants. This is illustrated in panels A and C of Supplementary Figures 1 and 2, where the 21J (Delta) variant sequences are divided into five and four distinct networks, respectively; in panels C, where Delta and Omicron sequences are separated; and in panel E, where 21L (Omicron) and 22C (Omicron) sequences are split into two distinct networks.

To explore the broader evolutionary relationships between these networks, phylogenetic analysis is the appropriate method, emphasizing the complementary nature of Minimum Spanning Networks (MSNs) and phylogeny. Phylogeny reconstructs evolutionary links between clearly distinct sequence groups, as demonstrated in the phylogenetic trees in panels B, D, and F of Supplementary Figures 1 and 2. These trees were generated using the same dataset as the MSNs for comparison. The trees in Supplementary Figure 1 were created using RAxML-version 8.2.12(4) with the GTRCAT substitution model and 100 bootstrap replicates, while those in Supplementary Figure 2 were produced with PAUP version 4.0a (build 168)(5) and 500 bootstrap replicates. Only nodes with support values of 75% or higher were kept in the consensus trees. As with MSNs, the background colors in the phylogenetic trees indicate the viral variant, following the same color scheme. The tree leaf colors correspond to the network nodes in the MSNs. MSNs offer a detailed view of evolutionary connections between closely related sequences, providing finer resolution than traditional phylogenetic methods. Conversely, phylogeny identifies evolutionary relationships between groups of genomically distinct sequences, even when prior knowledge is lacking (e.g., when analyzing a novel pathogen). However, phylogeny is most effective when applied to groups of genomically distant sequences. In panels B of

Supplementary Figures 1 and 2, phylogeny is unable to distinguish or relate some networks identified by VirNA because the sequences are too similar, which is where MSNs are more suitable.

VirNA can also detect multiple progenitors for a single sequence, suggesting potential cases of convergent evolution or recombination. This feature is inherent to Minimum Spanning Networks(6) and has been confirmed through the application of VirNA algorithm both to the simulated and real datasets.

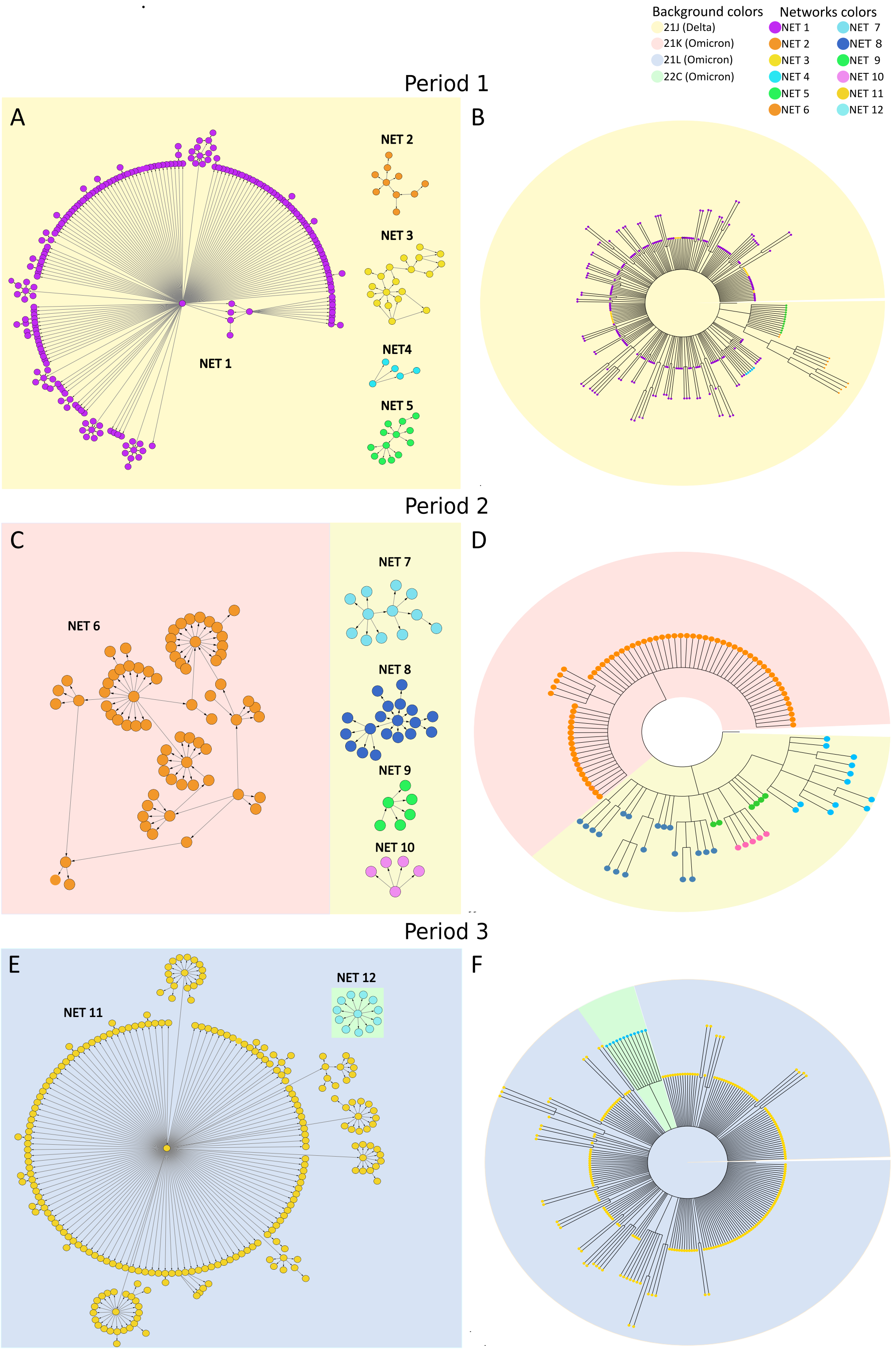


***Supplementary Figure 1**: MSNs and phylogenetic trees computed on multiple time periods. MSNs (panels A, C, and E) and phylogenetic trees (panels B, D, and F) computed by VirNA and PAUP(5) on the sequences from the periods 1, 2 and 3. The orientation of the arrows is determined by the haplotypes, following the principle of mutation accumulation. The color code used in the MSNs distinguishes the different viral clades and is maintained in the terminal nodes of the phylogenetic trees. The background color of the phylogenetic trees indicates the viral variant the terminal nodes belong to.*

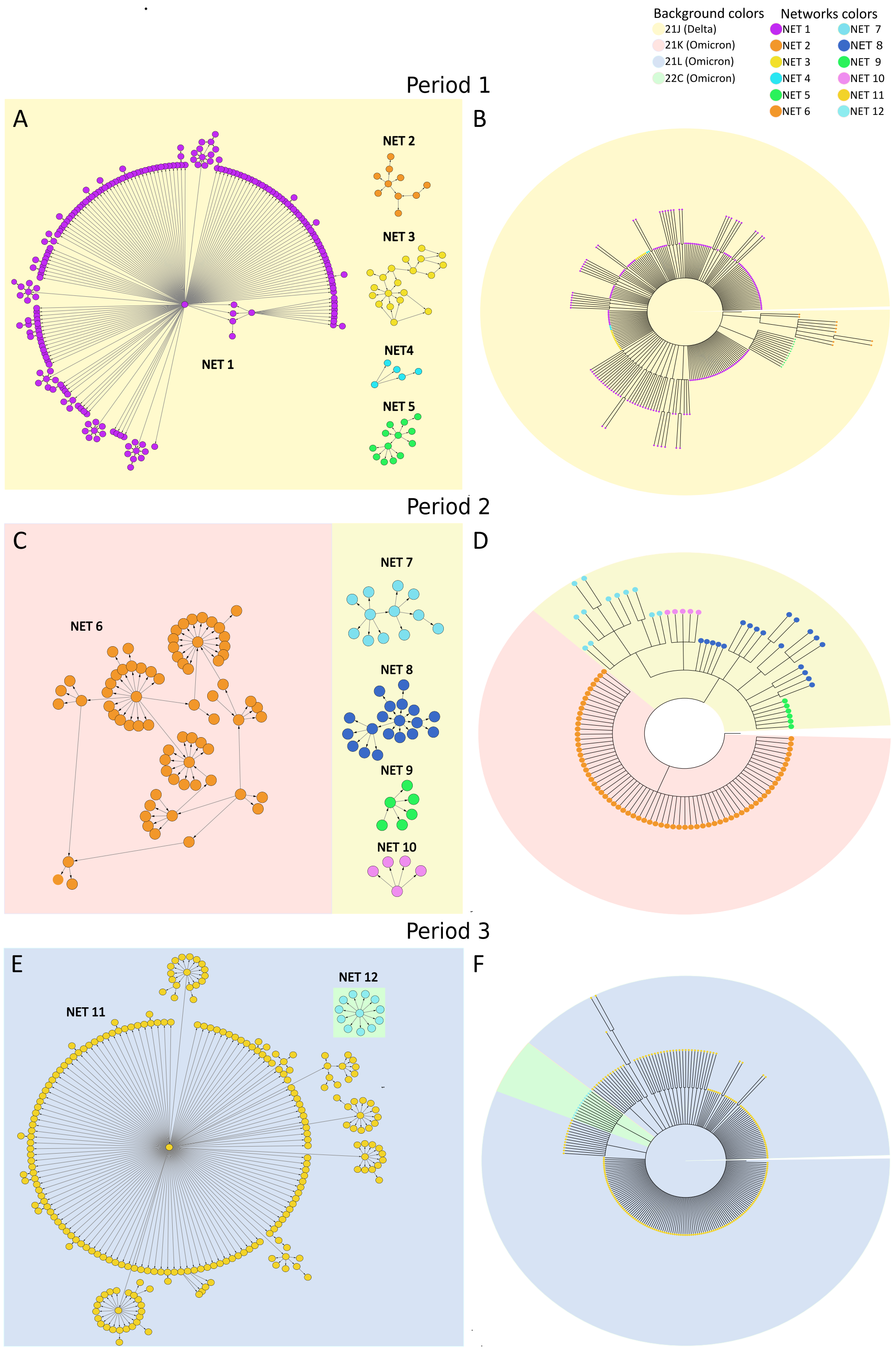


***Supplementary Figure 2****: MSNs and phylogenetic trees computed on multiple time periods. MSNs (panels A, C, and E) and phylogenetic trees (panels B, D, and F) computed by VirNA and RAxML(4) on the sequences from the periods 1, 2 and 3. The orientation of the arrows is determined by the haplotypes, following the principle of mutation accumulation. The color code used in the MSNs distinguishes the different viral clades and is maintained in the terminal nodes of the phylogenetic trees. The background color of the phylogenetic trees indicates the viral variant the terminal nodes belong to.*

### S2. MONKEYPOX CASE STUDY

The Monkeypox virus, which has seen a recent surge in global circulation, presents an abundance of genomic data making it another suitable candidate for testing VirNA. The initial dataset comprised 4707 genomic sequences obtained from the comprehensive EpiPox GISAID database(1) on January 23, 2023. The genomic sequences underwent a filtering process based on multiple criteria considering both genomic sequences and their associated metadata. The chosen sequences exhibit the following features: a complete collection date in YYYY-MM-DD format, a complete genome (>= 197000 b), absence of ambiguous characters, no head and/or tail gap trails or with gap trails shorter than 90 bases, and no insertions and/or deletions causing frameshift. The cleaned genomic sequences were aligned in an all VS all fashion along with their reference sequences (GenBank ON563414.3) using the Nextstrain augur tool(3) with default parameters. A refined dataset of 254 sequences was obtained, with the records globally sampled between May 14, 2022, and October 31, 2022. The manageable size of this dataset eliminated the need to select specific time periods. All the sequences belong to the IIb clade.

VirNA clusters the Monkeypox sequences into three distinct networks of non-compatible sequences, as shown in Supplementary Figures 3 and 4, panel A. The color code of the nodes is consistent across the MSNs and the phylogenetic tree. The tree in Supplementary Figure 3 was created using RAxML-version 8.2.12(4) with the GTRCAT substitution model and 100 bootstrap replicates, while the tree in Supplementary Figure 4 was produced with PAUP version 4.0a (build 168)(5) and 500 bootstrap replicates. Only nodes with support values of 75% or higher were kept in the consensus trees. The well-known limitations of phylogenetic algorithms in precisely positioning very similar genomic sequences within the tree are evident throughout the entire tree structure (Supplementary Figure 3, panel B), where a significant portion remains unsolved due to low bootstrap values, resulting in extensive polytomies. The networks generated by VirNA could suggest, in this specific case, the co-circulation of different viral strains, all belonging to the IIb viral variant. This implies that clade IIb may be further subdivided, as indicated by the three isolated clusters generated by VirNA.

Also in this case study VirNA demonstrates fine-grained resolution, revealing relationships among very similar haplotypes within a group, but also identifying co-circulating non-compatible haplotypes. However, phylogeny remains the preferred method for reconstructing a complete evolutionary history and understanding relationships among the isolated groups/clusters of sequences identified by VirNA. Thus, the Monkeypox case study affirms the complementary nature of these two methods, as also demonstrated in the SARS-CoV-2 case study.

VirNA can identify multiple putative progenitors for a single node in the Monkeypox dataset, as previously observed in SARS-CoV-2, a characteristic inherent to Minimum Spanning Networks(6). Due to the frequency of this phenomenon in Monkeypox dataset, we highlighted these occurences in the largest network generated (Network 1, panel A of Supplementary Figures 3 and 4). In these figures, the red nodes indicate entries with multiple ancestral nodes, along with the corresponding progenitors. The tables in panel C of Supplementary Figures 3 and 4 illustrate an exemplificative group of haplotypes responsible for generating the network, while the phylogenetic trees in panel B show how the phylogenetic algorithms positioned these sequences within the tree structure. For instance, nodes G and I are progenitors of node H, as indicated by the directional arrows and confirmed by analyzing the haplotypes in panel C. The mutations defining node H are a combination of the mutations from nodes G and I plus additional mutations, adhering to the compatibility requirement. However, the phylogenetic analysis struggles to accurately place these sequences within the tree structure. This challenge is evident in the phylogenetic trees in panels B of Supplementary Figures 3 and 4, where the sequences are frequently separated and part of large polytomies. An interesting aspect of this feature is linked to the directionality of the connections that follows a mutation accumulation pattern, leading to closed acyclic paths that ultimately form directed acyclic graphs, a specific feture of VirNA algorithm.

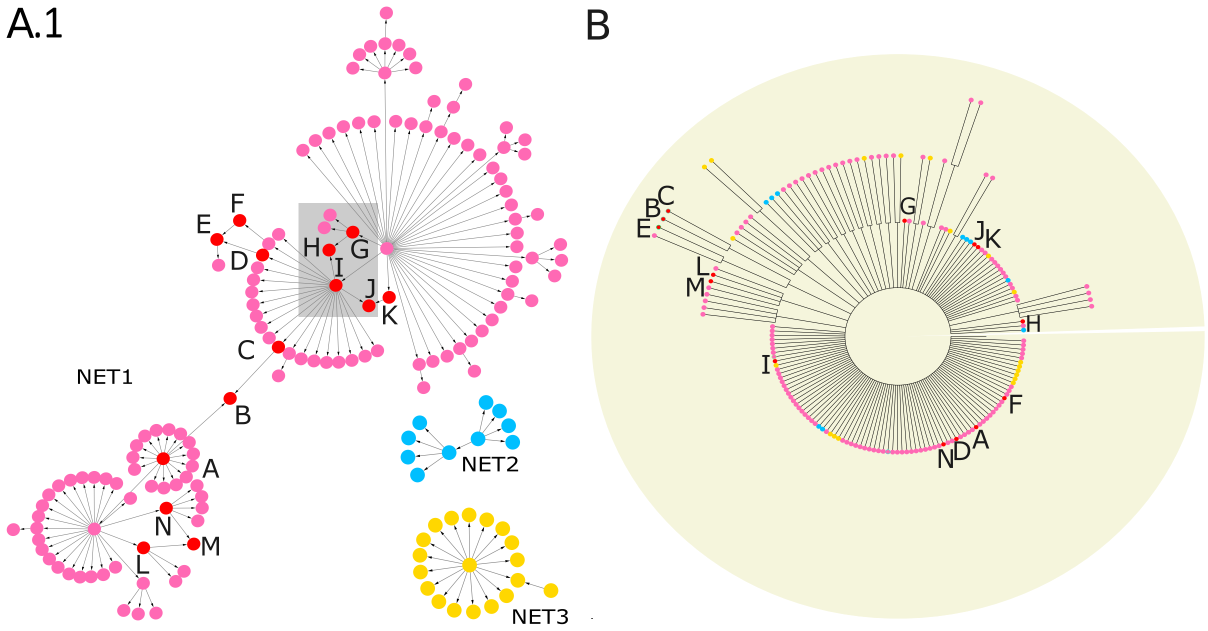


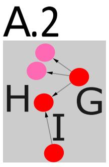


C

| Node | Set of mutations |
| --- | --- |
| G | G187050A |
| H | T1-, G187050A, C189276T, A198244- |
| I | A198244- |

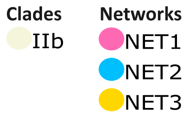


*Supplementary Figure 3: MSNs and phylogenetic tree computed on MonkeyPox sequences. MSNs computed by VirNA on IIb MonkeyPox sequences (A). The red nodes indicate the records with multiple ancestors along with their ancestors. The table indicates exemplificative haplotypes characterizing some of the underlined nodes (C). The phylogenetic tree, computed through RAxML(4), highlights where the nodes with multiple ancestors and their ancestors are located in the tree topology (B). The orientation of the arrows is based on the haplotype (from the pattern with less mutations to the pattern with more mutations). The color code of the MSNs highlights the networks identified for each Monkeypox group of non-compatible sequences by VirNA, with each network characterized by its own color. The nodes of the phylogenetic trees respect the same color code. The background of the phylogenetic tree indicates the viral variants the sequences belong to.*

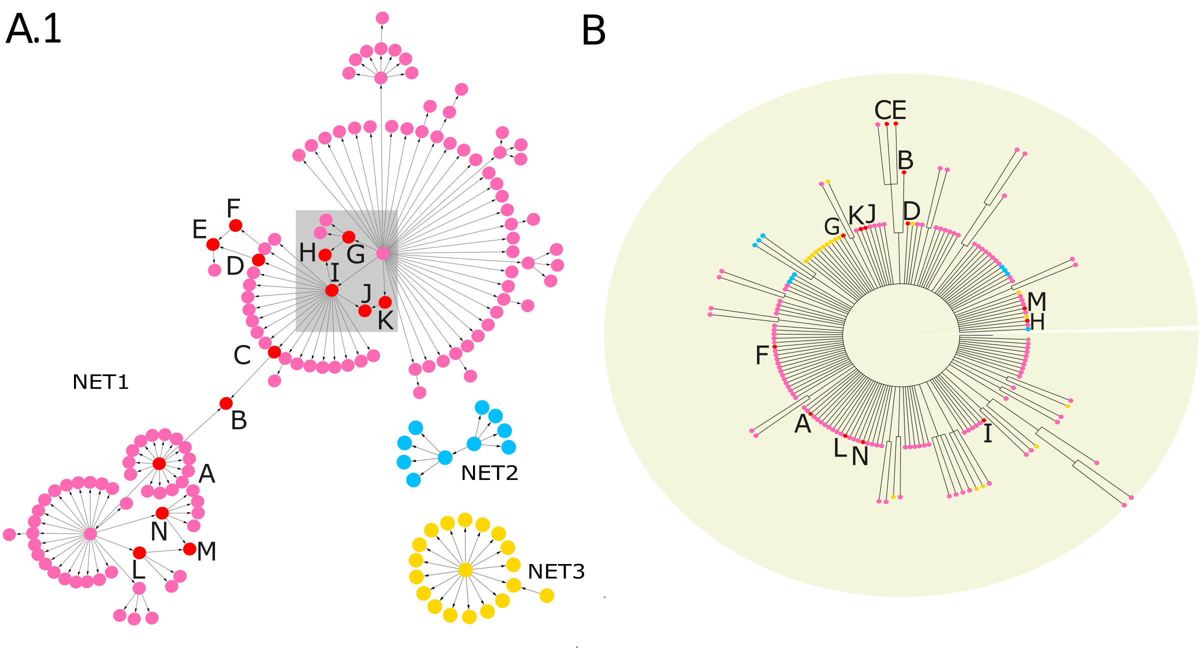


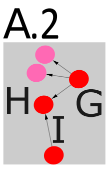


C

| Node | Set of mutations |
|---|---|
| G | G187050A |
| H | T1-, G187050A, C189276T, A198244- |
| I | A198244- |

Clades: IIb

Networks: NET1, NET2, NET3

*Supplementary Figure 4: MSNs and phylogenetic tree computed on MonkeyPox sequences. MSNs computed by VirNA on IIb MonkeyPox sequences (A). The red nodes indicate the records with multiple ancestors along with their ancestors. The table indicates exemplificative haplotypes characterizing some of the underlined nodes (C). The phylogenetic tree, computed through PAUP(5), highlights where the nodes with multiple ancestors and their ancestors are located in the tree topology (B). The orientation of the arrows is based on the haplotype (from the pattern with less mutations to the pattern with more mutations). The color code of the MSNs highlights the networks identified for each Monkeypox group of non-compatible sequences by VirNA, with each network characterized by its own color. The nodes of the phylogenetic trees respect the same color code. The background of the phylogenetic tree indicates the viral variants the sequences belong to.*

### S3. PERFORMANCE TEST OF VirNA

VirNA, Pegas(7) and PopArt(8) have been tested on five sets of SARS-CoV-2 sequences composed by 1000, 2000, 5000, 10000 and 15000 sequences respectively. VirNA is always faster then the other two tools as shown in Supplementary Table 1.

| **Sequence number** | **PopArt**(8) | **Pegas**(7) | **VirNA** |
| --- | --- | --- | --- |
| **1000** | 3 min | 3 min 25 sec | 52 sec |
| **2000** | 6 min | 1h 15 min 49 sec | 3 min 8 sec |
| **5000** | Data loading error | Network computation:<br>9 h 43 sec<br>Network drawing:<br>Pending and stopped after 3 days | 18 min 16 sec |
| **10000** | Data loading error | Not recorded: excessive computational time | 1 h 12 min 11 sec |
| **15000** | Data loading error | Not recorded: excessive computational time | 2h 44 min 43 sec |

***Supplementary Table 1: Performance test of VirNA.*** *Comparison of execution times in minutes of PopArt*(8), *Pegas*(7) *and VirNA.*
